# A quasi-analytic solution for real-time multi-exposure speckle imaging of tissue perfusion

**DOI:** 10.1101/2023.04.20.537736

**Authors:** Daniel A. Rivera, Chris B. Schaffer

## Abstract

Laser speckle contrast imaging (LSCI) is a widefield imaging technique that enables high spatiotemporal resolution measurement of blood flow. Laser coherence, optical aberrations, and static scattering effects restrict LSCI to relative and qualitative measurements. Multi-exposure speckle imaging (MESI) is a quantitative extension of LSCI that accounts for these factors but has been limited to post-acquisition analysis due to long data processing times. Here we propose and test a real-time quasi-analytic solution to fitting MESI data, using both simulated and real-world data from a mouse model of photothrombotic stroke. This rapid estimation of multi-exposure imaging (REMI) enables processing of full-frame MESI images at up to 8 Hz with negligible errors relative to time-intensive least-squares methods. REMI opens the door to real-time, quantitative measures of perfusion change using simple optical systems.

## 1. Introduction

Laser speckle contrast imaging (LSCI) is a relatively inexpensive widefield imaging method that allows for assessment of tissue perfusion and vascular flow across entire fields of view with high spatiotemporal resolution [1, 2]. LSCI gives a measure of flow based on speckle contrast, *K*, or the degree of blurring of a laser speckle pattern over a spatial region in a time-integrated image due to moving scatterers. Because of the simplicity of the technique, requiring only a laser, diffuser or beam expander, optical filter and camera, LSCI has been used extensively in neurophysiology for assessing perfusion and functional activity [3]. It has also gained some traction in clinical applications, such as in burn assessment, dermatology, and surgery [4], where it provides similar clinical capabilities to laser doppler flowmetry but with additional spatial information. LSCI, however, is limited to producing qualitative measurements of flow due to influences from the laser coherence, mismatches between the sensor pixel size and laser speckle size, and tissue scattering properties, which was addressed by the introduction of multi-exposure speckle imaging (MESI) [5-9]. In MESI, images with different exposure times are acquired and fit to a more robust model that accounts for laser coherence and mismatch between speckle size and pixel sensor size with the normalization parameter, β, for tissue scattering properties with the fraction of dynamic scattering parameter, ρ, and for directional vs. random scatterer motion by varying the light scattering model.

Some work has been done to bring quantitative MESI into clinical practice [10], and while the data acquisition is feasible in a clinical setting, the data analysis requires too much time to enable the real-time viewing needed for clinical application. The bottleneck for the use of whole field-of-view MESI in real-time applications is the need to fit speckle contrast values from multiple exposures at each pixel to models with the four parameters described above to reconstruct a single MESI image [11], a process that can take several minutes and up several hours depending on the image size and the machine used, which is why datasets are typically processed by restricting analysis to smaller, sparse regions of interest (ROI) over vessels or parenchymal tissue [12]. We find that processing a 1-megapixel image takes ∼20 minutes using 24 physical cores for parallel processing. To fit every image set of that size taken over the course of a minute at a 3-Hz acquisition rate would require 60 hours, making the use of ROIs for temporal analysis a necessity. Some methods have used calibrations before or during an imaging session to estimate the β and ρ terms to improve LSCI estimations [13] or speed up MESI fitting [10, 12, 14], but have the limitations of not accounting for spatial or temporal variability in β, caused by aberrations or out-of-focus surfaces or system fluctuations, or for temporal variability in ρ from, for example, changes in the fraction of tissue with flowing blood cells due to a stroke [11]. Other groups have incorporated machine learning to decrease the time required for processing MESI images [15, 16], although machine learning approaches require extensive training data and will likely be system specific.

Here we propose a quasi-analytic solution, noted as a Rapid Estimation of Multi-exposure Imaging (REMI), with a modification to incorporate spatial averaging of some parameters denoted as spatial-REMI (sREMI), to determine the scattering model, estimate the normalization and scattering parameters, and approximate the value of the correlation time with relatively high accuracy and up to 1500× faster than traditional least-squares fitting. The increase in speed allows for real-time assessments of perfusion for experimental or surgical applications after taking a few image sets and can be implemented with a rolling buffer of image data. We validate the accuracy of the method using simulated data, with and without artificial noise, and compare the performance to that of least-squares fitting for simple and mixed scattering models, demonstrate further improvements in accuracy and processing time by utilization of logarithmically-spaced exposure times over the typically-used exposure times, and finally compare the *in vivo* measurements taken from LSCI, least-squares fitting, and sREMI in a mouse model of photothrombotic stroke.

## 2. Theory of Laser Speckle Contrast Imaging

For LSCI, coherent light that has reflected off a rough, randomly scattering surface is imaged on a camera sensor, where the photons interfere constructively and destructively, producing the bright-dark spots of a speckle pattern. Moving scatterers, namely cells moving through blood vessels in tissue, introduce temporal fluctuations in this interference pattern which blurs the speckles within the image when integrated over a set exposure time. The degree of blurring increases with increasing speed of the moving scatterers. Speckle contrast, *K*, is quantified as

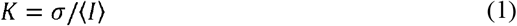

where σ is the standard deviation and ⟨*I*⟩ is the average of the pixel intensities calculated over a 5×5 or 7×7 window that is moved across the image (Fig. 1A). The challenge of speckle contrast imaging is then relating flow speed to this speckle contrast. The fluctuations in light intensity can be described by the electric field autocorrelation function, *g*_1_(τ), given by

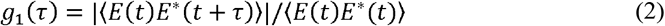

where *E* is the complex electric field at time *t*, and τ is the time delay. For single scattering from scatterers moving in random directions, g_1_(τ) follows a negative exponential, or Lorentzian, where

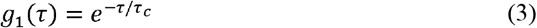

with τ_c_ as the correlation time. With faster moving scatterers, the speckle pattern will undergo more fluctuations and the autocorrelation function will decay sooner, resulting in a decreased correlation time.
The electric field cannot be easily measured to characterize g_1_(τ). The intensity, *I*, of the scattered light, however, is related to the electric field by *I* = *EE*^*^[1, 9] and can be measured by a camera sensor. The intensity autocorrelation function, g_2_(τ),

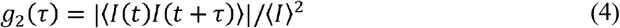

can be characterized and related to g_1_(τ) through a Siegert relation,

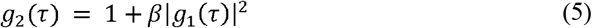

where β accounts for speckle averaging from the mismatch between the sizes of the laser speckles and sensor pixels [17, 18], as well as for imperfect light source coherence [19]. Speckle contrast was related to the correlation time by Bandyopadhyay et al. [20] by noting that the spatial variance in intensity, *σ*^2^, is related to the second moment of the intensity autocorrelation function such that

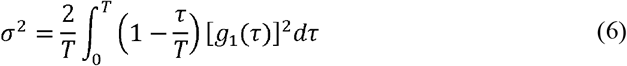

where *T* is the exposure time. In combining (1, (3, and (6, we get the relation between speckle contrast and the correlation time

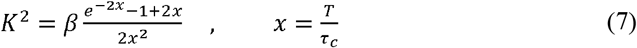

In LSCI, the β term can be determined by calibration to a static phantom. The use of a single exposure time, *T*, usually between 1 and 10 ms [21], allows for high temporal resolution imaging of relative blood flow speed. If the exposure is long enough compared to the anticipated correlation time, (7 can be simplified to estimate flow as

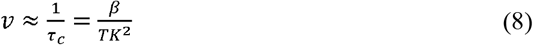

LSCI, however, fails to account for scattering by static components, which causes significant loss in sensitivity to moving scatterers [9, 13]. To address the limitations of LSCI, the autocorrelation function, g_2_(τ) was expanded by Parthasarathy et al [9] to include a scattering term from non-moving scatterers as

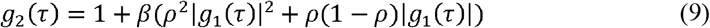

where ρ is the fraction of light that is scattered by moving scatterers (e.g. the fraction of dynamic scattering), given as 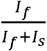 with *I*_*f*_ being the portion of fluctuating light, and *I*_*s*_ being the portion of statically scattered light. The resulting relation to speckle contrast is given by:

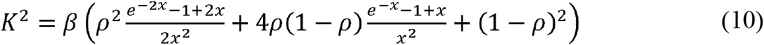

In order to fit the additional parameters, images are taken at multiple exposure times (Fig. 1B) with contrast values fit to (10 as a more quantitative measure in a method known as multi-exposure speckle imaging (MESI) [5-9]. The nonergodic noise normally added as a constant can be considered negligible in systems with low readout and sensor noise, or with additional noise reduction schemes [22].

**Fig. 1.**
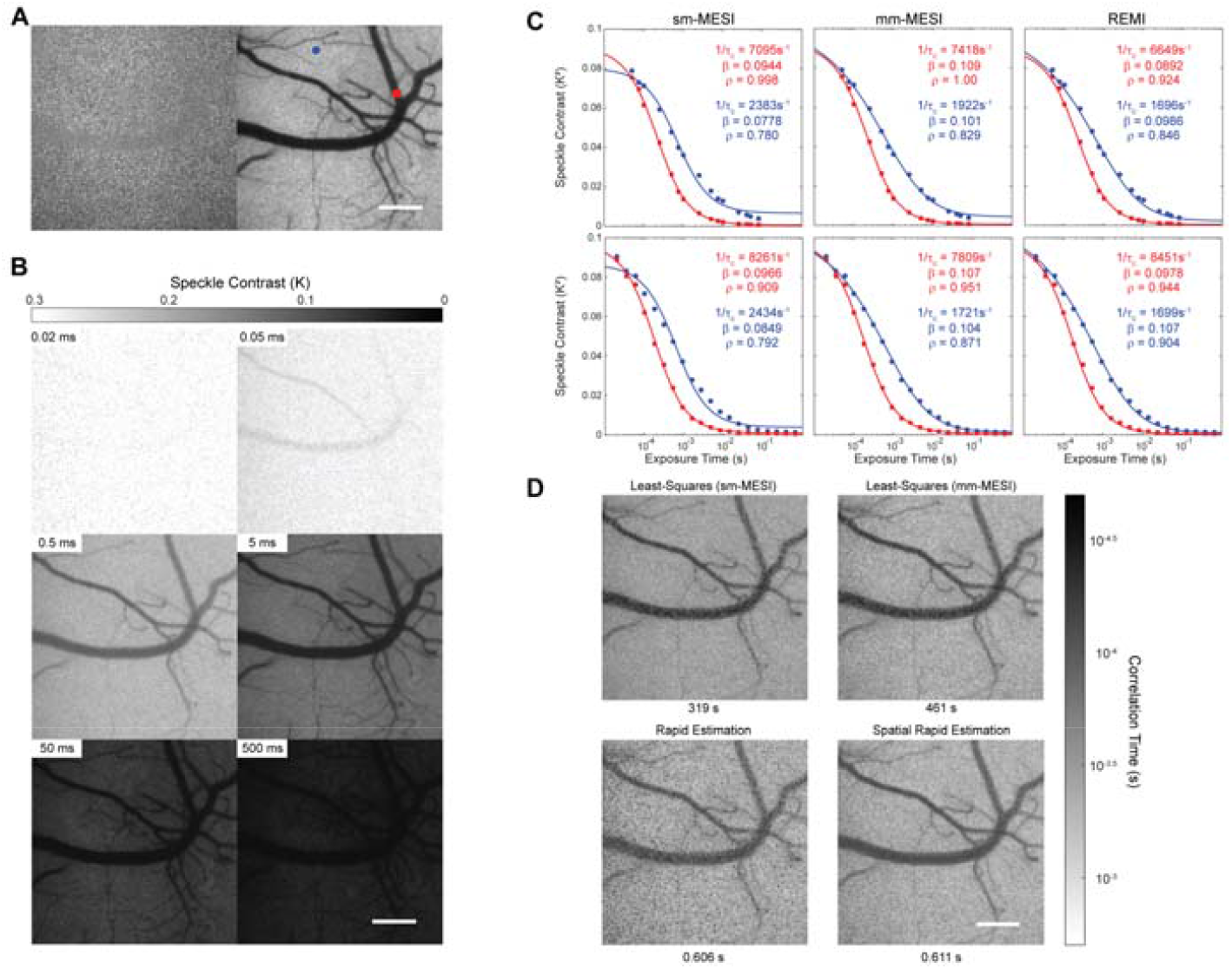
Laser speckle imaging and fitting examples. (A) Raw laser speckle image (5-ms exposure, left), and processed speckle contrast image (right). (B) Processed laser speckle images at varying exposure times from 20 μs up to 500 ms. Scale bar = 200 μm. (C) Different models are fitted to the speckle contrast data taken from indicated points in A, with the red point situated on top of a larger surface vessel and the blue point situated over a parenchymal region. The top row uses the exposure times commonly found in the literature [7, 26] from 50 μs – 80 ms, while the bottom row uses 20 logarithmicly-spaced exposure times from 20 μs – 650 ms, with all exposures listed in Table S1. Simple-model fitting (sm-MESI) shows multiple regions of overshoots and undershoots. Mixed-model fitting (mm-MESI) and Rapid MESI Estimation (REMI) follow the data more closely. (D) Example MESI images produced using simple-model fitting (top left), mixed-model fitting (top right), REMI (bottom left), and spatial-REMI (bottom, right). Images are 788 × 788 pixels. Required processing times are indicated below the images.

Work done by Postnov et. al [23] showed that while (10 worked well for the case of multiple scattered light from scatterers undergoing ordered movement, as seen in larger blood vessels, the model was less sensitive to flow changes in parenchymal tissue, where multiple scattering from scatterers undergoing unordered movement dominates [24]. Postnov, et al. thus modified g_1_(τ) to account for different scattering models [20, 24] with single or multiple scattering events, or for ordered or unordered movement of the scatterers. The electric field autocorrelation function was taken to be

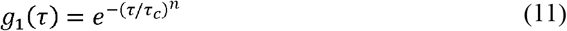

where *n* varies from 2 for single scattering from scatterers with ordered motion (single-ordered scattering; only observed in largest vessels, and not included in model here due to the photon mean free path length being significantly smaller than many vessels [24]), 1 for single-unordered (a small fraction of the detected light, as single scattering from deep-lying capillaries is rare [25]), or multiple-ordered scattering (dominant process for scattering from aligned flow of cells in arterioles and venules), and 0.5 for multiple-unordered scattering (dominant process for scattering from unaligned flow of cells in capillary networks in the brain parenchyma) (Fig. S1). In setting *n* = 0.5, the relation between speckle contrast from parenchymal regions and correlation time becomes

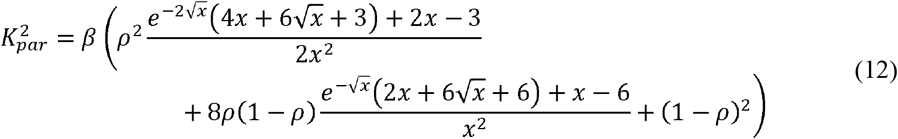

while the *n* = 1 expression for speckle contrast is now used just for vessels in which we rename (10 as 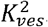. We use a mixed model combining (10 and (12 as

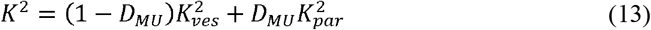

where *D*_*MU*_ is equal to 0 for pixels that contain only a vessel and 1 for pixels that contain only parenchyma. This mixed scattering model was shown to be far more sensitive to alterations in flow, for example in stroke models, as compared to the multiple-ordered, or single-unordered (*n*=1) scattering model [23]. In fact, we see that the fit to speckle contrast from parenchyma is improved over the traditional MESI fit using (10 alone (Fig. 1C). *D*_*MU*_ is allowed to have any value between 0 and 1 in the fits, accounting for pixels that contain variable fractions of larger vessels (ordered scattering) and parenchyma (unordered scattering).

## 3. Principles for rapid estimation of multi-exposure imaging (REMI)

Typically, regression methods seek to iteratively solve many variables simultaneously which requires many, often repeated, calculations. Our proposed method, Rapid Estimation of Multi-exposure Imaging (REMI), aims to solve for the unknown variables in a specific order using a set of limiting conditions from the model proposed in (13, drastically reducing the required number of calculations. We find REMI leads to fairly optimal fits and output values nearly matching more computationally intensive least-squares methods (Fig. 1C) with entire output images of similar clarity (Fig. 1D).

To begin, we consider three limiting conditions in (13: when the exposure time matches the correlation time (*T* = *τ*_*c*_), when the exposure time is much shorter than the correlation time(*T* < < *τ*_*c*_) or approaches 0, and when the exposure time is much greater than the correlation time (*T* > > *τ*_*c*_) or approaches ∞. For the condition where *T* = *τ*_*c*_, *x* is set to 1 in (10 and (12, which, combined in (13, gives a value, 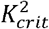, as:

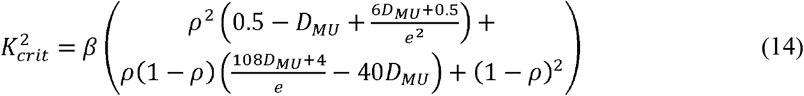

Using the latter two conditions, we get that the lim_*x*→0_ *K*^2^ → *β* and lim_x→∞_ *K*^2^ → *β* (1 − *ρ*)^2^ which, combined, allows us to solve for ρ as:

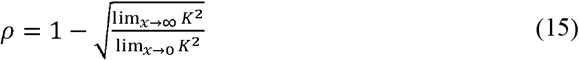

Obtaining these limiting conditions in practice is not feasible due to hardware or sacrifices to acquisition rate. Shorter exposure times are limited by available light or by camera hardware, meaning that extremely short exposures are not feasible for approximating β. On the other hand, extending the camera exposure to determine ρ would significantly reduce the resulting acquisition rate and introduce large amounts of ambient light and sensor noise. To address these physical limitations, we extrapolate from collected data using the first-order derivative with respect to *ln(T)*. In essence, we aim to estimate the remaining difference in contrast between the shortest and longest acquired image exposures and the extreme limiting conditions (Fig. 2A left, black arrows) by relating these differences to the first-order logarithmic derivative. The resulting derivatives of (10 and (12 yield:

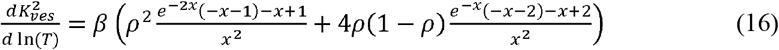

for multiple-ordered scattering, and

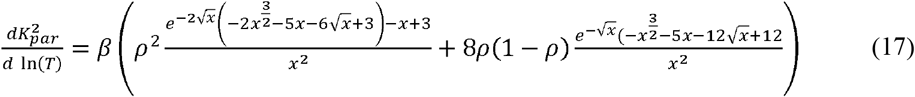

for multiple-unordered scattering and are represented in Fig. 2A, right.

**Fig. 2.**
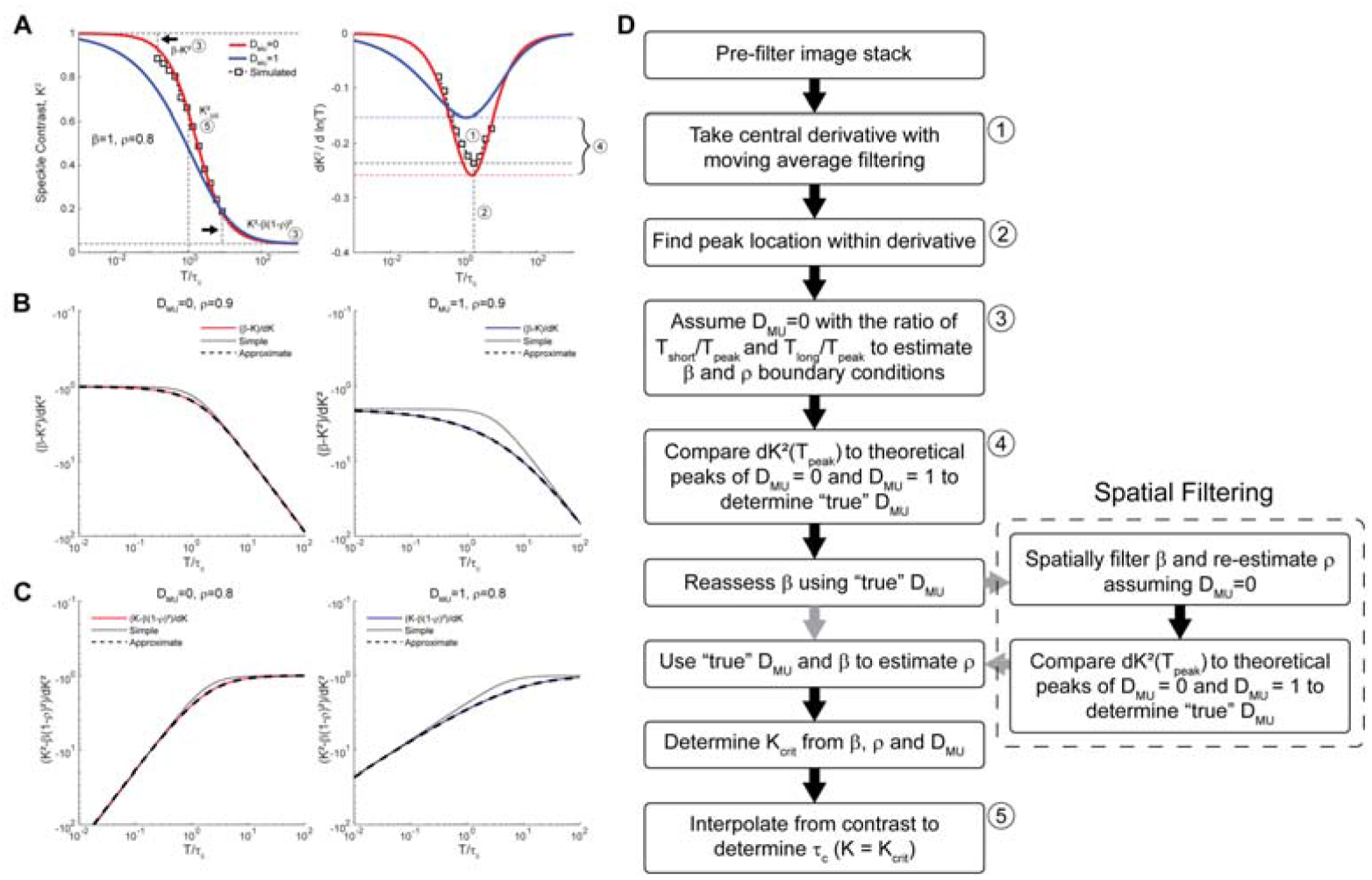
Underlying principles for implementing REMI. (A) Example plot of the ordered multiple scattering (red) and unordered multiple scattering (blue) models (left) and their respective derivatives (right) with example simulated data as white squares. (B) Ratio of *(K*^*2*^*(0)-K*^*2*^*(T))* to the derivative, *dK*^*2*^*(T)/d(ln(T))* (solid line) for extrapolation to determine the boundary condition setting β, with a simple lowpass fit (dotted line) and the expanded fit (dashed line). (C) Ratio of *(K*^*2*^*(T) – K*^*2*^(∞)) to the derivative, *dK*^*2*^*(T)/d(ln(T))* (solid line) for extrapolation to determine the boundary condition setting ρ, with a simple highpass fit (dotted line) and the expanded fit (dashed line). Solid lines are color-coded to match the scattering models from A. (D) Flowchart for the REMI algorithm. Numbered steps in the flowchart are presented in numbered bubbles in A. The gray arrows indicate the optional spatial filtering of β for sREMI.

When plotting the derivatives, we see noticeable differences in peak amplitude and location of the peak in the first-order derivative for each scattering model. While the peaks appear proximal to where *T* = τ_c_, the location for the minimum peak of *dK*^*2*^, noted as *T*_*peak*_, is shifted in relation to the true τ_c_ between ∼0.8τ_c_ and ∼3.3τ_c_ depending on the degree of dynamic scattering, ρ, and the scattering type, *D*_*MU*_ (Fig. S2A). Simply finding the peak of the differential will not give the true τ_c_ value due to the unknown shift induced by the unresolved parameters. To solve for the missing parameters, β and ρ, we examine the difference between *K*^*2*^(*T*) and the limit as *T* → 0 and *T* ∞ 0, whereby:

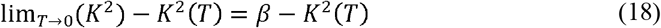

and

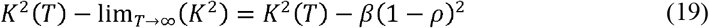

and take the ratio with respect to the slope at the same values of *T*, defined as 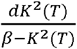 (Fig. 2B) and 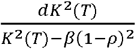 (Fig. 2C). When plotted in a log-log space, the former relation resembles a lowpass function while the latter resembles a highpass function, with both cases the curves. While a simple lowpass and highpass transfer function-like equation, such as 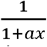 reaching a constant value (Fig. 2C, D) which is expected due to the asymptotic behavior of (dotted lines), can be applied to approximate this relationship, the empirical expansions provided by (20 and (21 give a more accurate approximation (Fig. 2B-C, dashed lines):

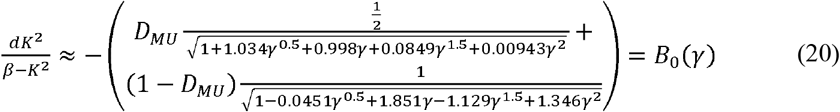

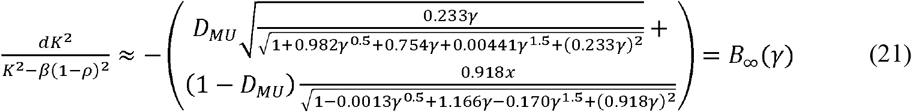

The produced decay curve shows a shift dependent on ρ when γ is set to equal *x* (Fig. S2B, C). We find that *T*_*peak*_ also shifts, such that when setting *γ*= *T/Y*_*max*_, the decay curves in (20 and (21 align across different values of ρ. It appears that the shifts in *T*_*peak*_ closely match those of the curves seen in Fig. S2B, C (although this cannot be confirmed analytically), making the use of the first-derivative peak location a reliable keystone for this approximation method. Rearranging (20 and (21 to solve for the limiting conditions gives:

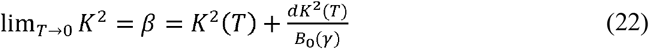

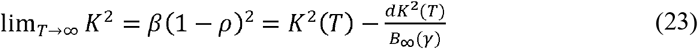

which allows us to estimate the boundary conditions with available data. Once the limiting conditions are known, 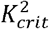 can be calculated using (14 and the correlation time of the dataset can be determined by simple interpolation.

## 4. Implementation of REMI

From (14, approximation of 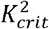 where *T* = τ_c_ requires estimation of the scattering model, *D*_*MU*_, the fraction of dynamic scattering, ρ, and the normalization factor, β. From (20 and (21, β and ρ can be estimated only when *D*_*MU*_ and *T*_*peak*_ are known. Of these, *T*_*peak*_ is the first variable that can be determined simply by locating the minimum peak location of *dK*^*2*^*/d(ln(T))* (Fig. 1B). We use *T*_*peak*_ in (22 and (23 to approximate β and ρ assuming *D*_*MU*_ = 0, then compare the minimum amplitude of *dK*^*2*^*/d(ln(T))* to the theoretical minimum of (16 and (17 using the estimated terms. *D*_*MU*_ is then set to a value between 0 and 1, determined by the distance between *dK*^*2*^*(T*_*peak*_*)/d(ln(T))* and the theoretical amplitudes for the two scattering models (Fig. 2A right, curly bracket). Once the scattering model is determined, the limiting conditions of lim_*x*→0_ *K*^2^ and lim_*x*→∞_ *K*^2^ are estimated by using (22 and (23, now with the appropriate *D*_*MU*_. β is set to lim_*x*→0_ K^2^and ρ is calculated using (15. At this point, the scattering model, *D*_*MU*_, the fraction of dynamic scattering, ρ, and the normalization constant, β, have been solved. These parameters enable the use of (14 to set the 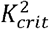 value which equals (*K*^2^(*T*) when *T* = τ_c,_, allowing for simple interpolation to solve. We provide more details on some of the critical steps in this process in the rest of this section.

### 4.1 Pre-processing discrete data

Laser speckle image data can be noisy, especially with fewer averaged speckle contrast images per exposure. Differentiating noisy discrete data only further exacerbates the noise. To reduce noise, the discrete data needs to be filtered in some way, but filtering can often suppress important higher frequency features. To balance this, we use a 1:4:1 weighted moving average filter for evenly-spaced exposures in the logarithmic domain, or a weighted nonuniform moving average for unequally-spaced typical exposure times [26] (see Table S1; see Supplementary Materials for details on nonuniform moving average filter). The central differential of the filtered data is then taken over intervals of *ln(T)* to achieve a semilogarithmic derivative. To further reduce noise, the derivative is filtered again using a simple 3-point moving average filter for equally spaced exposure times or a weighted-moving average filter for unequally spaced data with a width of 1 decade. Unequally spaced data is resampled using the filtered differential to interpolate and match the location of the exposure time of the central differential for further operations.

### 4.2 Determining the scattering model and the unsolved parameters

To determine the scattering model, we first need to find the steepest point of the slope of the speckle contrast, *K*^*2*^, to determine *T*_*peak*_. The exposure time values of the most negative value of *dK*^*2*^*/d(ln(T))* and the closest two surrounding points are used in a quadratic Newton interpolation polynomial while setting the derivative of the polynomial to 0 to locate *T*_*peak*_. *dK*^*2*^*(T*_*peak*_*)/d(ln(T))* is determined by setting *T*_*peak*_ as the interpolation point of the polynomial. As a moving average filter is expected to attenuate the peak of the differential, we increase the amplitude by 5%, determined empirically, to compensate in the next 2 steps. Using the ratio of *T*_*peak*_ to the shortest and longest exposure times, we apply (22 and (23, assuming *D*_*MU*_ = 0 to approximate *K*^2^(0) and *K*^2^(∞), respectively. (15 is then used to solve for ρ.

To determine the correct scattering model, we compare *dK*^*2*^*(T*_*peak*_*)/d(ln(T))* to the theoretical max negative slope for both *D*_*MU*_ conditions. Repeatedly solving for these minimum values in (16 and (17 for comparison is computationally expensive. However, the peak value, when holding *D*_*MU*_ constant, only depends on ρ and is scaled by β. This simplification allows for empirically computing the peak values for both *D*_*MU*_ = 0 and *D*_*MU*_ = 1 as a function of ρ from 0 to 1 using the fminbnd() function in MATLAB and fitting a polynomial (this paper used an 10^th^ order polynomial; see Supplementary Materials for polynomial expression) for reference. In this way, we can now pass an M × N matrix of ρ values and output two M × N matrices of peak values for comparison with low computational overhead. We predict β and ρ using (16 and (17 while letting *D*_*MU*_ = 0 and calculate the theoretical peak amplitudes for *D*_*MU*_ = 0 and *D*_*MU*_ = 1 using ρ and β with the polynomial fits. The scattering model *D*_*MU*_ is then given as the distance of *dK*^*2*^*(T*_*peak*_*)/d(ln(T))* from the theoretical peak in the *D*_*MU*_ = 0 condition divided by the distance between the theoretical peaks for the *D*_*MU*_ = 0 and *D*_*MU*_ = 1 conditions, bounded between 0 and 1 (Fig. 2A; *right, bubble 4*).

### 4.3 Spatially averaging β for spatial-REMI (sREMI)

Up to this point, all data is processed in a pixel-by-pixel manner, incorporating no spatial information, which can result in significant salt-and-pepper noise (Fig. 1D, bottom-left). The normalization constant, β, is expected to vary slowly across the field-of-view due to optical aberrations and defocus across the imaging field. Therefore, β can be spatially filtered to reduce salt-and-pepper artifacts (Fig. 1D, bottom-right). (22 is used with the first estimate of *D*_*MU*_ to calculate an initial β. A gaussian filter with a width of at least 10% of the image field is then applied to the estimated β. We then re-estimate *D*_*MU*_ as outlined above, using the spatially filtered *K*^2^ (0) when initially setting *D*_*MU*_ = 0 (Fig. 2D, dashed outline).

### 4.4 Interpolation of the correlation time, τ_*c*_

Once the scattering model has been determined, we re-evaluate ρ using (23, setting *D*_*MU*_ at the inferred value. The “true” β and ρ are applied in (14 to determine the 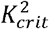 values for which *T* = τ_*c*_. For strong static scattering regions, 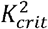 is taken as the minimum of 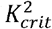 and 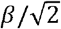 to prevent falsely high flow rate measurements due to a near-zero slope produced by very low dynamic scattering (ρ < 0.2). (Empirically, this limit was only reached with completely static scatters, such as a piece of paper.) We run a simple linear interpolation using two points that bound 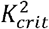 to get our final estimate of the correlation time. The first bounding point takes the the index, *idx*, of the last speckle contrast value above 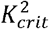 for each pixel. The next index, *idx*+1, then acts as second bounding coordinate for the interpolation:

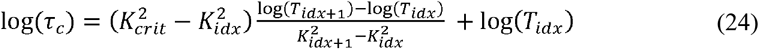

If the longest exposure remains above 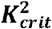, the last two indices are used for the interpolation. On the other hand, if the first value is already below 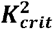, the first two indices are used.

## 5. Results

### 5.1 Performance on simulated data

Mixed-model least-squares fitting (e.g. to (13), simple-model least-squares fitting (e.g. to (10), REMI, and sREMI were applied to simulated data using both multiple-ordered scattering and multiple-unordered scattering models in MATLAB 2021a on an Intel® Xeon® CPU E5-2687W v4 @ 3.00GHz 24-core system running Ubuntu OS using the traditional exposure times and log-spaced simulated exposure times from Table S1. The parallel processing toolbox with 24 workers was utilized only for the least-squares fitting methods. In the simulated datasets, 1/τ_c_ was varied from 10^0^-10^5^ s^-1^ and ρ was varied from 0.5-1, with and without applying artificial uniform noise of 5% of the true value. The correlation times determined from each method is plotted against the true simulated values in Fig. 3, with the impact of artificial noise shown in Fig. S3. Output parameters for β and ρ are plotted against the true 1/τ_c_ values for each method in Fig. S4. Least-squares fitting of the mixed model was highly accurate over a broad range from the shortest exposure time up until the 2^nd^ longest exposure time (Fig. 3A, S3A), and fit well for simulated data from both scattering models. As expected, the simple-model fit well for multiple-ordered scattering data (e.g. as we would find in a vessel), even beyond the shortest exposure time. However the accuracy for this model was low with multiple-unordered scattering data (e.g. as we would find in a parenchymal region) (Fig. 3B, S3B). REMI had high accuracy in estimating inverse correlation times between 10^2^ and 10^4^ s^-1^ for the exposures between 50 μs and 80 ms which aligned with the 3^rd^ shortest and 5^th^ longest exposures used (Fig. 3C, S3C) with improved performance seen using logarithmic exposures (Fig. 3D, S3D). By applying a spatial Gaussian filter (size = 500^2^ pixels, σ=250 pixels) to set β, the accuracy of sREMI was extended to the shortest exposure time and to the next longest exposure time (Fig. 3E, S3E). The spatial map used for testing sREMI is displayed in Fig. S3F.

**Fig. 3.**
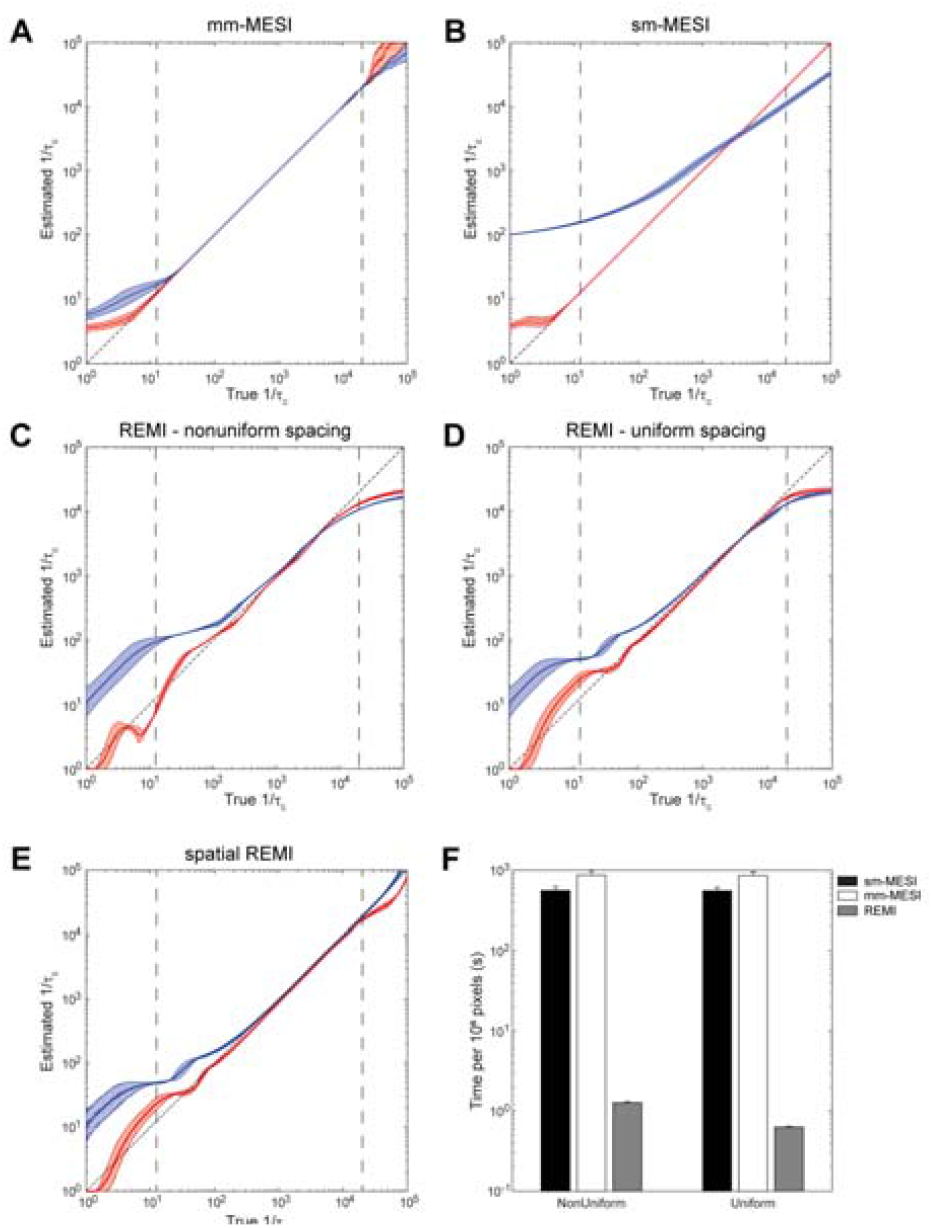
Performance of processing methods on simulated data. Plots of estimated correlation times vs. true correlation times for (A) mixed-model fitting, (B) simple-model fitting, (C) REMI using typical exposure times, (D) REMI using logarithmically spaced exposure times, and (E) sREMI using logarithmically spaced exposure times. Red traces correspond to multiple-ordered scattering data; blue traces correspond to multiple-unordered scattering data. Shaded regions represent standard deviations from log-space values. Vertical dashed lines in (A-E) represent the bounds set by the exposure times. (F) Bar plot indicates the time required to process 10^6^ pixels. Error bars represent variance across three repeats on each set of simulated data (clean-ordered scattering, clean unordered scattering, noisy-ordered scattering, noisy-unordered scattering).

The mixed-model least-squares fitting method had a slightly higher accuracy and maintained that accuracy over a broader range of correlation times, as compared to REMI, but took 700-1400× longer to process with 10^6^ pixels and 15 exposures (Fig. 3F). When varying the number of pixels and number of exposures, the least-squares fitting for both models showed a near linear dependence on the number of pixels with a small influence by the number of exposures while REMI was equally dependent on the number of pixels and exposure times. With evenly, logarithmically spaced exposure times, and using the compiled filtering convolution function, the exponent for the dependence on the number of exposures was further reduced to ∼0.5. The resulting correlation coefficients between true and estimated τ_*c*_ and the respective processing time coefficients for each method are given in Table 1.

**Table 1.** Correlation coefficients between the true τ_c_ and the estimated τ_c_ for each method and the empirical equation for required processing time.

| Method | R<sup>2</sup> (clean) | | | R<sup>2</sup> (noisy) | | | Processing Time<br>= $A N_{\text{pix}}^B N_{\text{exp}}^C$ | | |
| --- | --- | --- | --- | --- | --- | --- | --- | --- | --- |
|  | Ves | Par | Mix | Ves | Par | Mix | A | B | C |
| sm-MESI | 1.0000 | 0.9866 | 0.9326 | 0.9966 | 0.9771 | 0.9273 | 1.73e-3 | 0.899 | 0.048 |
| mm-MESI | 1.0000 | 1.0000 | 1.0000 | 0.9949 | 0.9861 | 0.9881 | 1.70e-3 | 0.924 | 0.092 |
| REMI | 0.9807 | 0.9790 | 0.9165 | 0.9699 | 0.9075 | 0.8767 | 9.06e-9 | 1.112 | 1.240 |
| REMI-log | 0.9934 | 0.9821 | 0.9149 | 0.9542 | 0.6942 | 0.8018 | 3.26e-8 | 1.106 | 0.542 |
| sREMI-log | 0.9923 | 0.9695 | 0.9188 | 0.9539 | 0.6465 | 0.7786 | 4.55e-9 | 1.271 | 0.442 |

Performance of each analysis method on the additional parameters, β and ρ, is demonstrated in Fig. S4. For REMI, we note an underestimation of β as the true correlation time falls near or below the lower range of exposures, which demonstrates the inability to accurately capture the rising slope of the correlation function (e.g. Fig. 2A). With sREMI, the additional spatial averaging strongly reduces the error in β. For the scattering coefficient, ρ, accuracy is impacted on two fronts. The first results from the same challenge as with the β term, in that the falling slope of the correlation function is not well constrained with the given exposures. The second impact is due to the underestimation of β, which by (15, inevitably influences the estimation of ρ. The former underestimation is to be expected as having low contrast at short exposure times can limit fits to small β values. Likewise longer correlation times, from slower moving scatterers, can be mistaken as nonmoving scatterers, resulting in a lower dynamic scattering fraction. Nonetheless, REMI and, especially, sREMI produce nearly as accurate an estimate of β and ρ as with the full mixed-model least-squares approach, with slightly higher errors in β in regions with high flow speeds (with REMI, but not sREMI) and slightly higher errors in ρ in regions with low flow speeds (with both REMI and sREMI).

### 5.1 Comparison between methods in a mouse model of photothrombotic stroke

To test the utility of REMI *in vivo*, a mouse was imaged under anesthesia with respiratory rate and heart rate monitoring by a piezoelectric sensor [27] while inducing a photothrombotic occlusion in an arteriole using Rose-Bengal solution and illumination by a green diode laser with full-width half-max diameter of ∼500 μm at the brain surface (Fig. 4A) [28-30]. The different processing methods were applied to image data taken from the mouse during repeated photothrombotic strokes. We used an initial exposure of 60 s at an average power of 1 mW, which induced a partial, transient occlusion, then waited some time before inducing a full occlusion by increasing the average green laser power to 2 mW. Average maps before and after photothrombotic occlusions with overlaid ROIs were produced using 100 frames from sREMI outputs (Fig. 4B,C). Movies of 3-point moving average frames produced using sREMI with a spatial Gaussian filter (size = 101^2^ pixels, σ = 50 pixels) on a 5-point moving average of speckle contrast images are shown in Supplementary Video 1 (partial occlusion) and Supplementary Video 2 (subsequent, full occlusion) with whole field-of-view maps of correlation times (τ_c_, left), normalization parameter (β, top-right), dynamic scattering fraction (ρ, middle-right), and the scattering model (D_MU_, bottom-right). Dynamic scattering is seen to decrease slightly during the first, mild photothrombotic stroke (Supplementary Video 1) with the second occlusion produced a significant decrease in dynamic scattering (Fig. 4B, C panel iii) and a notable transition between scattering models, due to decreased contribution from multiple, ordered scattering after the surface arteriole was fully clotted (Fig. 4B, C panel iv) (Supplementary Video 2).

**Figure 4.**
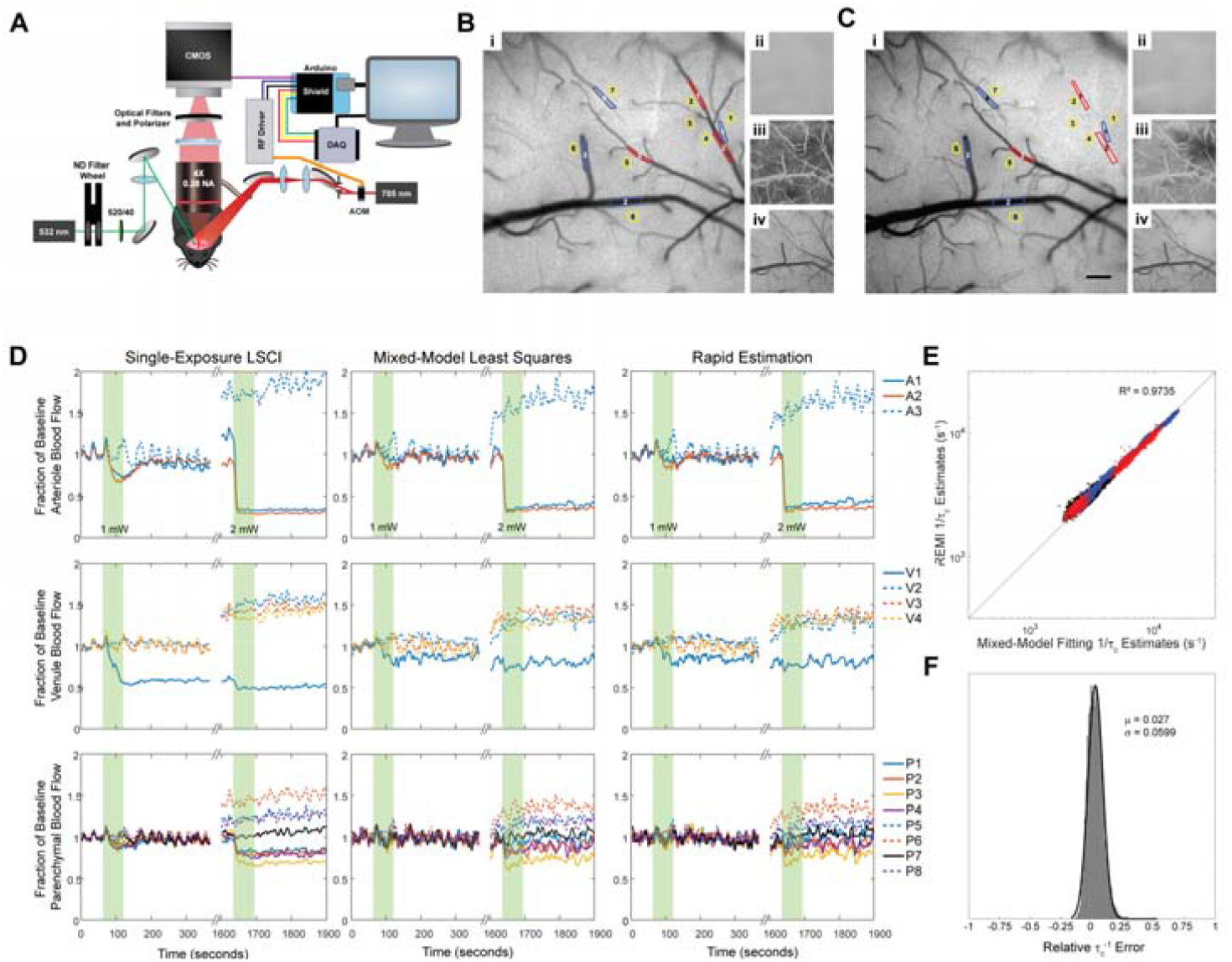
Processing of multi-exposure speckle images from a mouse model of photothrombotic stroke. (A) Simplified schematic of multi-exposure speckle imaging setup. A microcontroller controls pulse duration and transmitted laser power through the acousto-optic modulator (AOM) as well as camera exposure times for speckle imaging. The data acquisition (DAQ) board records signals for post-imaging synchronization. Neutral density (ND) filters control the power of the green laser used for producing the photothrombotic stroke and optical cleanup filters extinguish residual infrared light from the green laser to prevent interference with the laser speckle imaging. Maps produced by sREMI before (B) and after (C) photothrombotic stroke, showing correlation time (i), normalization coefficient (ii), dynamic scattering (iii), and scattering type (iv) with selected ROIs over arterioles (red), venules (blue), and parenchyma (yellow) indicated on the correlation time maps. Scale bar is 200 μm. (C) Traces of changes in perfusion from indicated ROIs in arterioles (top), venules (middle), and parenchyma (bottom) as measured using LSCI (left), mixed-model least-squares fitting (middle), and sREMI (right) during photothrombotic stroke (green bars) using a 10-point moving average to minimize fluctuations from heart rate. (D) Scatter plot of inverse correlation times from ROIs placed over arteriole (red), venule (blue), and parenchymal (black) regions between sREMI and mixed-model fitting. (E) Distribution of relative errors from sREMI measurements with mixed-model fitting as ground truth.

All methods were able to capture flow changes during the transient occlusion with 1 mW and following the full occlusion at 2 mW in the targeted arteriole (Fig. 4D, S5A), but the estimated magnitude of the flow decrease varied by method. After the 1-mW irradiation, there was a noticeable, transient drop in flow in the targeted arteriole that returned to baseline in a couple minutes (Fig. 4D, S5A top), indicating a partial occlusion that was released upon termination of the light exposure. Mixed-model fitting and sREMI estimated the flow speed decrease in the arteriole to be 17% and 18%, respectively, while simple-model fitting estimated a 15% reduction and LSCI estimated the drop to be ∼41%. With 2 mW of green light exposure, mixed-model fitting and sREMI measured a reduction of 68% and 66%, respectively, in the targeted arteriole from the initial baseline. Simple-model fitting showed a decrease of 60% while LSCI indicated a reduction of 72% from the initial baseline.

To quantify the degree of agreement between methods, we examined the correlation between the inverse correlation times across models, taking the mixed-model least-squares results as ground truth. The simple-model fitting had the overall highest correlation coefficient to the mixed-model fits (R^2^ = 0.9823, Fig. S5B), but there were large relative errors, with a bimodal distribution: 3.9 ± 2.9% and 20 ± 7.4% for the two peaks, respectively (Fig. S5C). In comparison to the mixed-model, the simple-model fitting tended to overestimate flow magnitude in parenchymal regions and in occluded vasculature, leading to the peak at ∼20% error (Fig. S5A). LSCI estimation of flow (Fig. 4D left column) tended to be significantly slower than the mixed-model for all ROI types, and had the lowest correlation (R^2^ = 0.7648, Fig. S5D, E) and largest errors relative to the mixed-model, again with a bimodal distribution: -52 ± 10% and -76 ± 5.6%. The errors are largely due to the inability of LSCI to account for changes in the fraction of light scattering that comes from moving scatterers and not accounting for a shift toward unordered scattering at the site of a vessel occlusion (Fig. 4B, C panel iii). We find that a reduction in the dynamic scattering term by ∼5%, as we saw during the mild stroke, results in a ∼20% underestimation of flow by LSCI, while not allowing the scattering model to change from more ordered to more unordered scattering, which was observed after the full occlusion at the location of the targeted vessel, results in an additional ∼15% reduction in the LSCI flow estimate. sREMI showed a high degree of correlation with the mixed-model least-squares fit in flow changes for all types of ROIs (R^2^ = 0.9785, Fig. 4E), and had the lowest relative error of 2.6 ± 6%, with a normal distribution for this error (Fig. 4F).

## 6. Discussion

In this paper, we have devised an algorithm that allows for real-time processing of multi-exposure speckle images without the need for least-squares fitting. We validate the efficacy of the algorithm on simulated data, further improve performance using uniform logarithmic exposure times compared to typically used exposure times, and accurately track changes in a mouse during photothrombotic stroke while using mixed-model least-squares fitting as a ground truth.

While REMI is shown to be much quicker than the least-squares fitting approaches, it comes with the drawback of being more influenced by signal noise. As seen in Fig. 1D, there is prominent salt-and-pepper noise present due to some pixels having higher variability in speckle contrast at the shorter exposure times. By applying a spatial filter to β, due to the slow variations expected across the field of view, the high variability at the shortest exposures is eliminated and thus a significant portion of the salt-and-pepper noise is removed (Fig. 1D). Averaging multiple subsequent REMI or sREMI output images further reduced fluctuations to appear as in Fig. 4B, C.

Unlike least-squares regression methods, REMI allows for estimation of all pixels in an image in a near-real time, or real-time, manner. The time required for REMI to operate scales near-linearly with the pixels and exposures when taken non-logarithmically spaced data, such that a 512×512×15 matrix takes ∼0.25 s to process. A major limitation in speed comes from the nonuniform moving average filter applied to the original data as well as the first-order differential using the uncompiled processing in MATLAB. We improved speed by equally spacing the exposure times on a logarithmic scale and applying a lower-level convolution function to filter the data, which reduces the processing time to ∼0.12 s. The time difference between sREMI and REMI is near-negligible, influenced primarily by the size of the Gaussian filter for spatially filtering β.

As REMI calculations are done across all pixels in parallel, the algorithm is slowed by memory allocations for large matrices associated with larger images, meaning that least-squares fitting methods would eventually take less time to run, though this only begins to occur at unreasonable image sizes on the order of megapixel to gigapixel side lengths. Of course, the image size for nonlinear least-squares regression methods to match the speed of REMI will also change with available parallel processing capabilities.

While single-exposure LSCI is a fast and simple alternative to the quantitative multi-exposure speckle imaging approach, it comes with the drawback of not being able to account for changes in static scattering, causing significant errors in flow measurements. During the mild photothrombotic stroke, flow reductions were estimated by LSCI to be much larger than with any other method. Based on Monte-Carlo simulations, the laser speckle signal integrates scattering signals from up to 700 μm below the tissue surface [25]. As the green laser light did not occlude the surface arteriole and flow speed in the arteriole returned to baseline following termination of the green laser light illumination, it is possible that the initial photothrombotic occlusions were restricted to the capillaries just beneath the surface, leading to fewer moving and more static scatterers but leaving the deeper-lying capillary network largely unaffected. The result would be a decrease in the dynamic scattering coefficient while holding the correlation time near constant or only slightly reduced, leading to errors in LSCI flow speed estimation. The sREMI approach applied to this data tracked these changes and produced estimates that closely matched the mixed model, least squares approach.

## 7. Conclusion

This paper demonstrates a method for quasi-analytically approximating the correlation times for MESI images in a rapid and reliable manner, with close agreement with the output from least-squares approximations, without the need for pixel-by-pixel curve fitting, removing a significant bottleneck for real-time viewing. The up to 1500× reduction in processing time allows for monitoring of blood flow changes in real time in clinical and research settings without extensive equipment or machine learning. With proper parallelization, an image set can be recorded, while converting a previous set to speckle contrast images and processing to MESI images then have all outputs displayed with no noticeable delays. All MATLAB code for the implementation of REMI and sREMI, as well as the example data from Fig. 1, is available on GitHub at: https://github.com/sn-lab/Speckle-REMI.

## Acknowledgements

We would like to thank Robert Hawkins for assisting with the Rose-Bengal stroke experiments and Dominick Romano for early assistance with the change of variables derivation process. We are grateful to David Boas for comments on an earlier draft of this manuscript.

