## Supplemental materials for "A quasi-analytic solution for real-time multi-exposure speckle imaging of tissue perfusion"

Methods

Testing on simulated data

The proposed algorithm was run on simulated data with standard exposure times as well as logarithmically-spaced exposure times between 50 µs and 80 ms. For the simulated data, ρ was uniformly varied between 0.5 and 1, τ_c_ was uniformly varied between 10^-5^ and 10^0^, and β was held constant at 0.10 while using both *D_MU_*=0 and *D_MU_*=1 conditions. Datasets were left clean or with added uniform noise of ±5% of the true pixel value and comprised of 10^6^ samples for each condition (8 × 10^6^ samples total). For assessing the effectiveness of the spatial filtering of β, a spatially varying dataset was produced for which τ_c_ ranged between 10^-5^ and 10^0^ (Fig. S3F), ρ was randomly set between 0.5 and 1, and *D_MU_* was randomly set as 0 or 1. Mixed-model nonlinear least-squares curve fitting used initial estimates of β=0.12, ρ=0.9, τ_c_=0.001, and D_MU_=0.5. Simple-model nonlinear least-squares curve fitting used the same conditions with the exclusion of D_MU_. To avoid issues with the random seed for introducing noise, the datasets were pre-generated to compare between REMI and nonlinear least-squares fitting. To test the dependence on the size of the dataset, random datasets were generated ranging from 64^2^-2048^2^ pixels with 10-25 logarithmically spaced exposures. Image processing was done in MATLAB 2021a on an Intel® Xeon® CPU E5-2687W v4 @ 3.00GHz 24-core system running Ubuntu OS. Least-squares fitting was performed using the parallel processing toolbox with 24 workers to match the number of physical cores. The use of REMI and sREMI was done without parallel processing or GPUs.

Animal preparation

All animal procedures were approved by the Cornell Institutional Animal Care and Use Committee (protocol 2015-0029) and were performed under the guidance of the Cornell Center for Animal Resources and Education. Adult C57BL/6 mice (n=2) at 3 and 9 months of age were used.

Mice were induced under anesthesia using 3% isoflurane in 100% oxygen, moved to a stereotactic frame, and maintained at 1-2% isoflurane during surgical procedures. Body temperature was regulated with a feedback-controlled DC temperature controller and heating pad (40-90-8D, FHC). The fur on the scalp was shaved and the area cleaned with alternating iodine and 70% ethanol solutions. Prior to any incisions, mice received ketoprofen (5 mg/kg, 2.0 mg/mL) and dexamethasone (0.2 mg/kg, 0.1 mg/mL) to reduce inflammation from the removal of the skull. 100 µL of bupivacaine (0.1% solution) was administered into the incision site as a local anesthetic. The mice received ~5-mm diameter craniotomies between lambda and bregma using a 0.7-mm dental drill, followed by placement of an 8-mm cover glass. Sterile saline was added to the surface of the brain before securing the cover glass to the dried skull surrounding the craniotomy with cyanoacrylate (Loctite). Dental cement sealed the craniotomy to restore intracranial pressure and maintain sterility. One mouse was allowed 3 weeks for recovery before imaging. A photothrombotic stroke was induced acutely in the second mouse.

Multi-exposure imaging setup

Multi-exposure speckle imaging was performed using a custom-built system (Fig. 4A). Illumination was provided using a 785-nm 300-mW laser diode (LD785-SEV300, ThorLabs). An acousto-optic modulator (AOMO 3100-125, Gooch & Housego) driven with an RF driver (1110AF-AEFO-1.5, Gooch & Housego) adjusted the amount of laser light that was diffracted to the animal so that the average image intensity remained the same for all the exposure times. Speckle images were taken through a 4× 0.28 NA objective (XLFluor, Olympus) onto a CMOS camera (acA2040-90umNIR, Basler). A linear polarizer and bandpass filter (769/41 nm, AVR Optics) were placed to enhance speckle contrast and reduce light from external sources, respectively. An Arduino Uno microcontroller communicated with a custom program written in MATLAB to set the diffracted laser amplitude and gate the pulses of light and camera acquisition using a specialized circuit board. Because of camera limitations, for exposure times shorter than 50 µs, the camera was triggered for 50 µs while the laser light was pulsed for the shorter exposure time. A DAQ board (USB-6001, National Instruments) recorded the TTL signals to the camera, the output AOM voltage, and the output of a custom animal sensor composed of a piezoelectric disk and filtering circuitry on which the animal is placed for monitoring respiratory rate and heart rate. Exposure times used for each experiment are listed in Table S1.

Table S1. Exposure times used in each experiment and simulation

| Experimental Protocol | Exposures (ms) |
| --- | --- |
| Standard exposures | 0.05, 0.075, 0.10, 0.25, 0.50, 0.75, 1.0, 2.5, 5.0, 7.5, 10, 25, 40, 50, 80 |
| Simulated log-spaced exposures | 0.05, 0.085, 0.14, 0.24, 0.41, 0.70, 1.2, 2.0, 3.4, 5.7, 9.7, 16, 28, 47, 80 |
| Extended exposures | 0.02, 0.035, 0.06, 0.10, 0.18, 0.31, 0.53, 0.92, 1.6, 2.7, 4.7, 8.2, 14, 24, 42, 73, 130, 220, 380, 650 |
| Photothrombosis exposures | 0.02, 0.036, 0.065, 0.12, 0.21, 0.39, 0.70, 1.3, 2.3, 4.1, 7.5, 14, 24, 44, 80 |

Multi-exposure speckle imaging

For imaging, the mice were induced under 3% isoflurane in 20% oxygen, placed in a stereotactic frame, and maintained at 1-2% isoflurane while monitoring respiratory and heart rate via the piezoelectric sensor. Body temperature was regulated with a feedback-controlled heating pad. For the first mouse, 30 images for each of the standard exposure times were taken at 12-bit resolution in a cyclical manner. The raw images were processed into speckle contrast images using a 7×7 window, then averaged into a single 2000×2000×15 speckle contrast matrix. An additional 10 speckle images were averaged for each of the 20 extended exposure times between 20 µs and 650 ms with the same resolution and bit-depth to produce a 2000×2000×20 speckle contrast matrix. For the induced photothrombotic stroke, the photothrombotic set of exposures in Table 1 at were taken at ~1.5-2 Hz for at least 5 minutes.

Induction of a photothrombotic stroke

For induction of a photothrombotic stroke, we used a Rose Bengal solution which photochemically produces reactive oxygen species in the presence of light, with a continuous-wave green laser diode (λ = 532 nm) (Compass 215M-50, Coherent) as the illumination source. A series of neutral density filters reduced the power to ~1-2 mW at the imaging plane and a 520/40 nm optical filter (ThorLabs) removed residual NIR light from the green laser so as not to interfere with the speckle imaging. A 200-mm focal length lens focused the beam down to ~500-µm diameter while a periscope steered the beam to a target location (Fig. 4A). The green beam was placed over an arteriole using a laser speckle contrast map of the vasculature as reference. Once positioned, the beam was blocked before retro-orbitally injecting the mouse with 50 µL of Rose-Bengal solution (10 mg/mL). Image acquisition began 30-60 s prior to unblocking the green diode laser. The green laser remained unblocked for 60 s with the times of illumination onset and termination marked on the DAQ board by a short TTL pulse.

Comparing the change in blood flow following photothrombotic stroke across each method

For processing of the photothrombotic stroke, speckle images were averaged in a 5-point rolling average before processing through each method. 4-ms exposures from the datasets taken before and after induction of the stroke were used to select ROIs around the affected arteriole. For least-squares fitting of the simple-model and mixed-model, only pixels in the ROIs were evaluated to reduce the total processing time required. REMI values were processed in a whole image manner using the spatial averaging for β with the average values taken from the ROIs. LSCI values were approximated with 4-ms exposure images using Eq. 8 with β values determined by mixed-model fitting (we note this is a more involved estimation of β than would typically be used for LSCI, where data from a completely static sample, such as a piece of paper, is more commonly used).

Nonuniform moving average filter

For unequally-spaced data in the logarithmic domain typically used in the literature, a nonuniform moving average filter with a total width of 1 decade, or 0.5 decades across the center point, was applied to reduce system measurement noise. The points within range are weighted by the distance to the center timepoint as $wt= (\frac{w}{2}-\left| T-T_{center} \right|)$ and the filter applied as

$$K_{f}^{2}\left( T_{center} \right)=\frac{\sum_{i}^{I} K_{i}^{2}wt_{i}}{\sum_{i}^{I} wt_{i}}$$

(S1)

Polynomial expansions

Table S2 shows the coefficients for the 10th order polynomial expansion of the speckle contrast derivative relative to ρ, which is used to reduce computational overhead in determining the scattering model in Step 4 from Fig. 2.

Table S2. Polynomial expansion of *dK^2^(T_peak_)/d(ln(T)) as a function of ρ*

| Multiple-Ordered/Single-Unordered | |  | Multiple-Unordered | |
| --- | --- | --- | --- | --- |
| $\boldsymbol{x}^{\boldsymbol{n}}$ | $\boldsymbol{c}_{\boldsymbol{n}}$ |  | $\boldsymbol{x}^{\boldsymbol{n}}$ | $\boldsymbol{c}_{\boldsymbol{n}}$ |
| 10 | -0.020475343735539 |  | 10 | 0.053297852491085 |
| 9 | -0.000249703516762 |  | 9 | -0.267813703593650 |
| 8 | 0.193138590579442 |  | 8 | 0.529732554893485 |
| 7 | -0.360479187939359 |  | 7 | -0.522690178203463 |
| 6 | 0.282977235939114 |  | 6 | 0.299320834277943 |
| 5 | -0.159691928427907 |  | 5 | -0.130682424041211 |
| 4 | 0.008759628227216 |  | 4 | -0.002907089625296 |
| 3 | -0.042800145778791 |  | 3 | -0.039727987048522 |
| 2 | 0.740574034436267 |  | 2 | 0.463936419535563 |
| 1 | -1.283491167540141 |  | 1 | -0.764925668142888 |
| 0 | 0.000000130185034 |  | 0 | 0.000000051719825 |

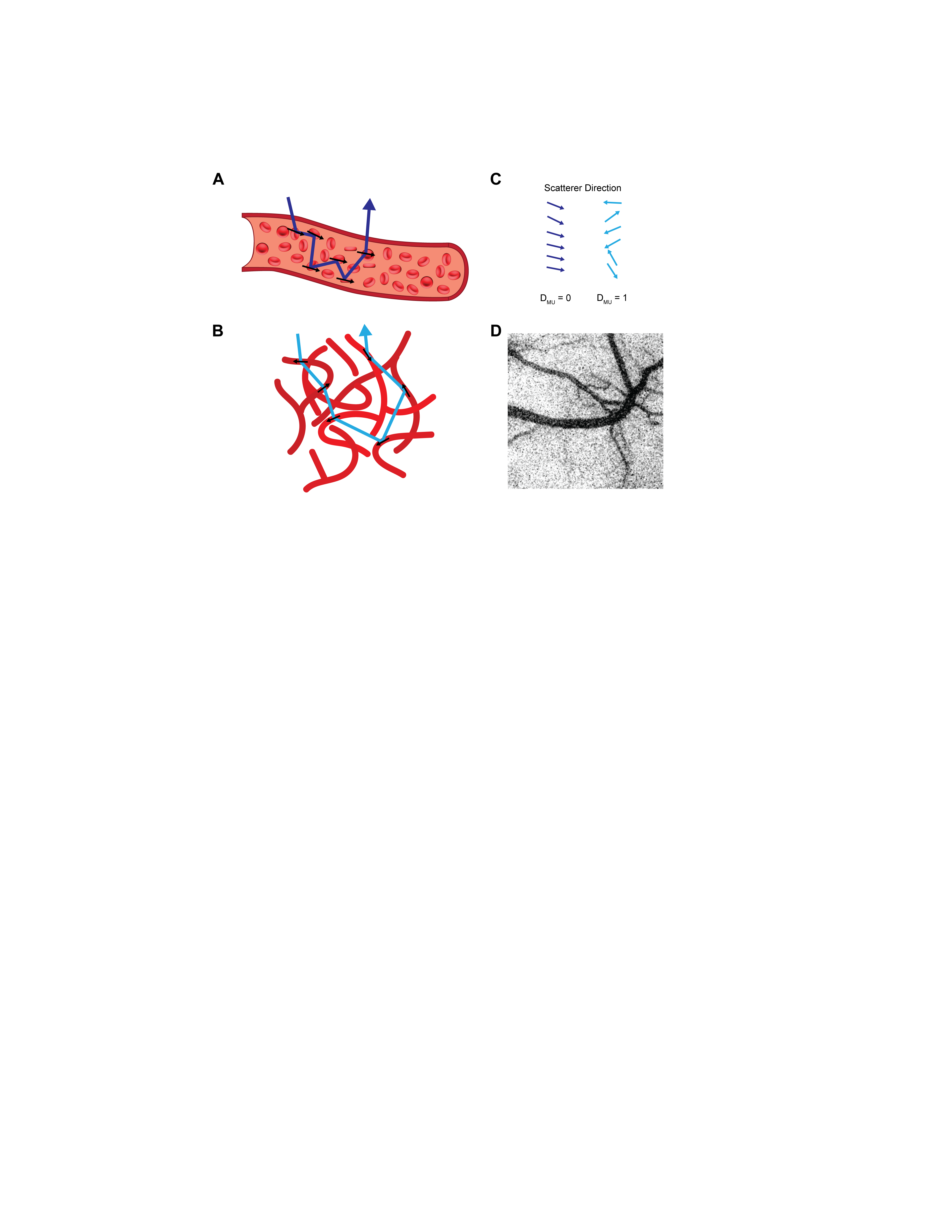

Figure S1. Schematic of the multiple scattering models for vascular and parenchymal regions. (A) Model vessel representing multiple scattering of a photon on scatterers moving in an ordered direction. (B) Example of a capillary bed with photons scattering off cells moving in random directions. (C) Arrows representing the scatterers’ direction of movement and indicating the appropriate scattering model to capture the dynamics. Colors correspond to the photon path color in A and B. (D) Scattering model coefficient of a processed multi-exposure image using sREMI. Vessels are near zero (black), with a gradient moving outward from the vessels to parenchyma, where the coefficient approaches one (white).

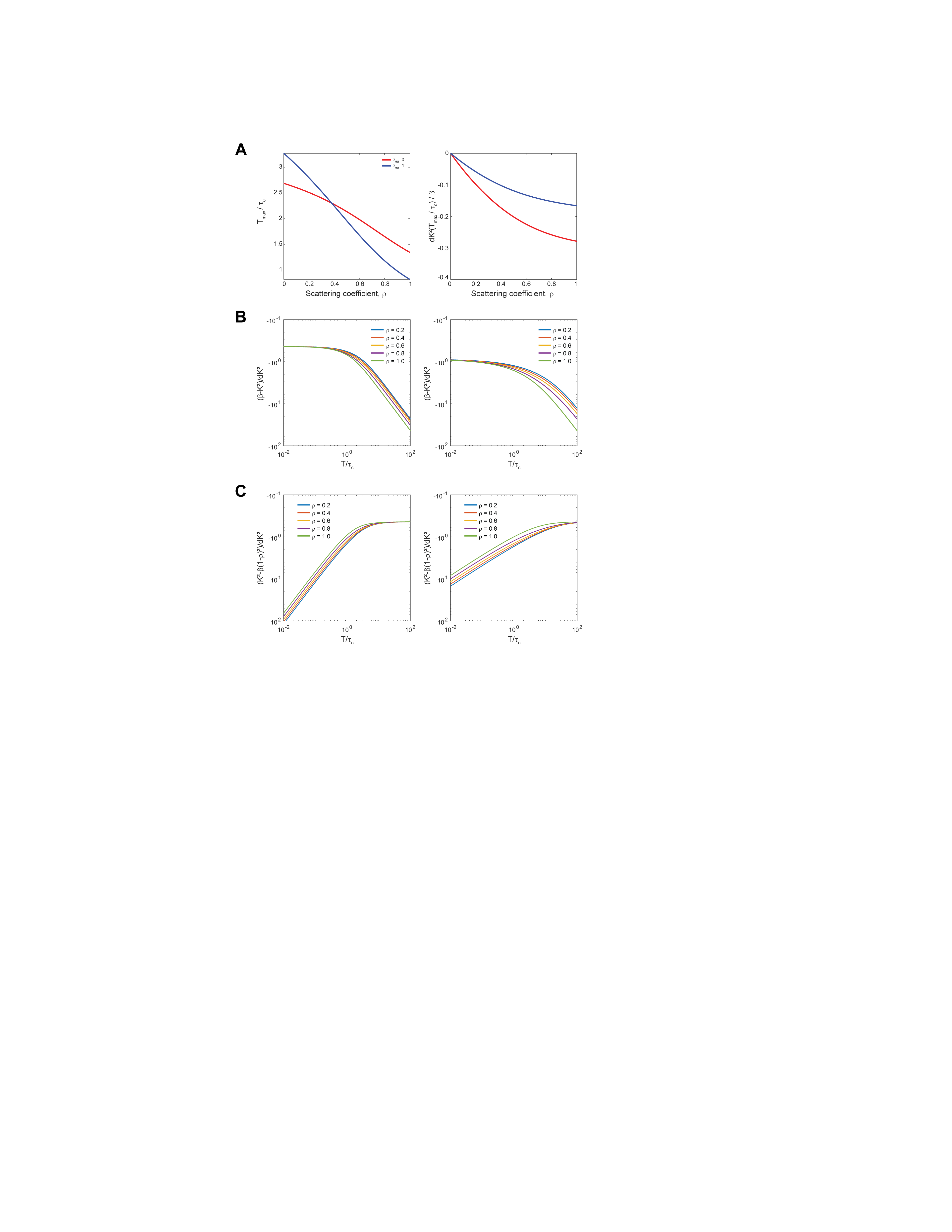

Figure S2. Influence of dynamic scattering on function behavior. (A) Shift in differential peak location (left) and peak amplitude (right) as a function of ρ for multiple-ordered scattering (red) and multiple-unordered scattering models (blue). (B) Shift in ratio of *(K^2^(0)-K^2^(T))* to *dK^2^(T)/d(ln(T))* with respect to τ_c_ for various ρ for multiple-ordered scattering (left) and multiple-unordered scattering (right). (C) Shift in ratio of *(K^2^(T)-K^2^(∞))* to *dK^2^(T)/d(ln(T))* with respect to τ_c_ for various ρ for multiple-ordered scattering (left) and multiple-unordered scattering (right).

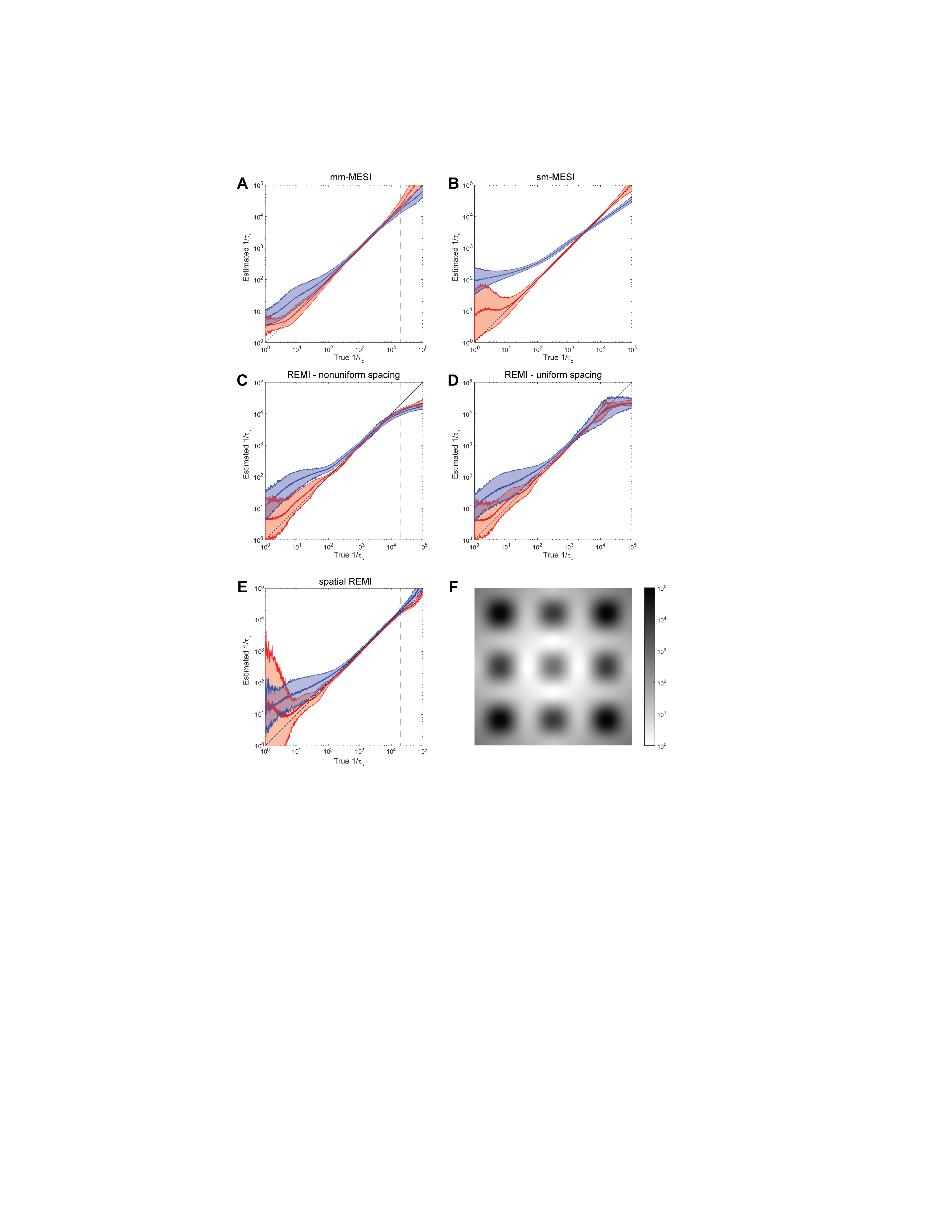

Figure S3. Impact of artificial noise on correlation time measurements. Plots of true τ_c_ to estimated τ_c_ values for (A) mixed-model fitting, (B) simple-model fitting, (C) REMI, (D) log-spaced REMI, and (E) sREMI (log-spaced), all with 5% added variability in speckle contrast values. Traces tend to follow the *y = x* line, though the deviation from the line due to noise varies by method. (F) Spatial distribution of inverse correlation times (s^-1^) used in sREMI test set.

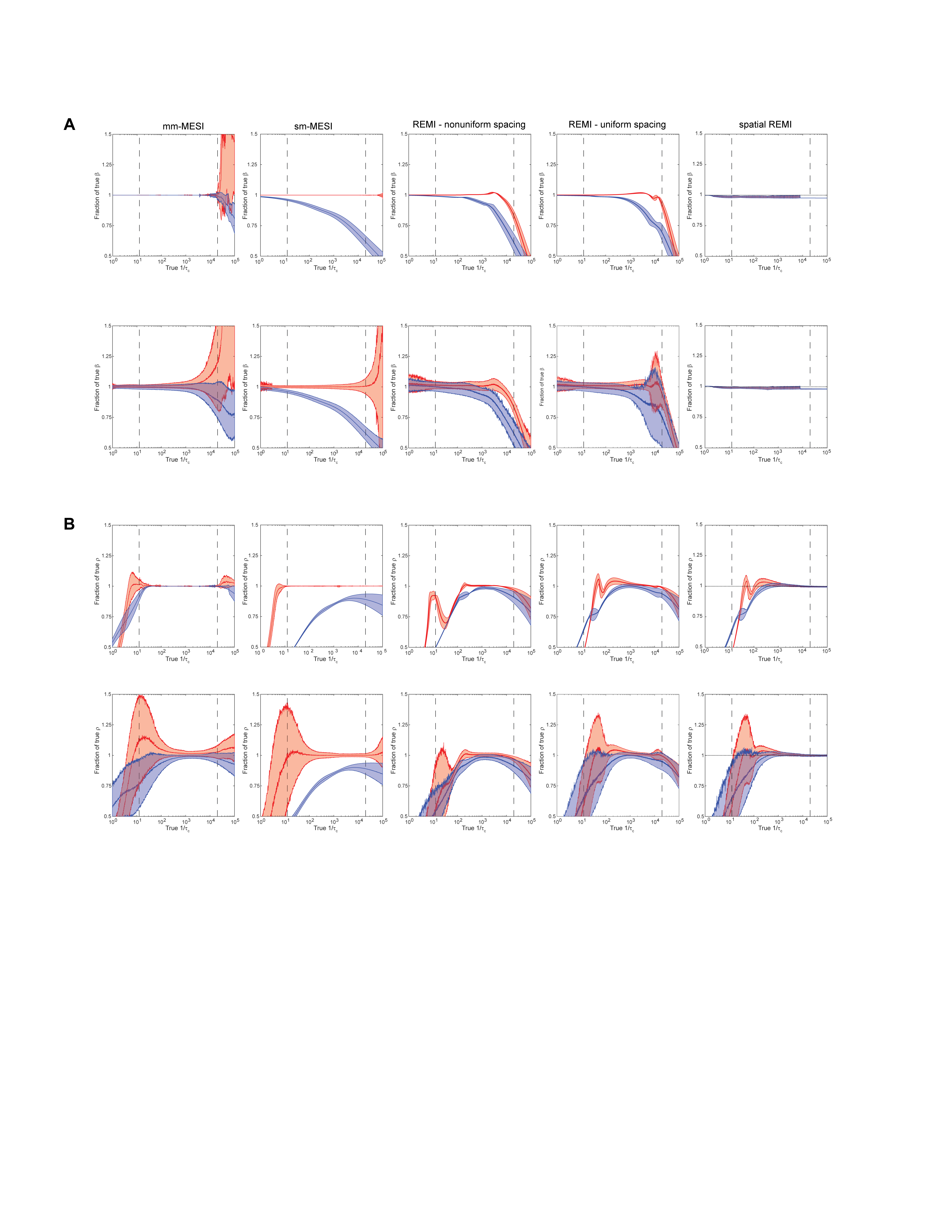

Figure S4. Performance of β and ρ parameters for each processing method. Ratio of estimated to true (A) β and (B) ρ as τ_c_ varies with clean (top) and noisy (bottom; 5% variability in speckle contrast) simulated data for mixed-model fitting, simple-model fitting, REMI, log-spaced REMI, and sREMI (log-spaced) from left to right. Red traces represent multiple-ordered scattering data and blue traces represent multiple-unordered scattering data. Shaded regions represent one standard deviation from the mean.

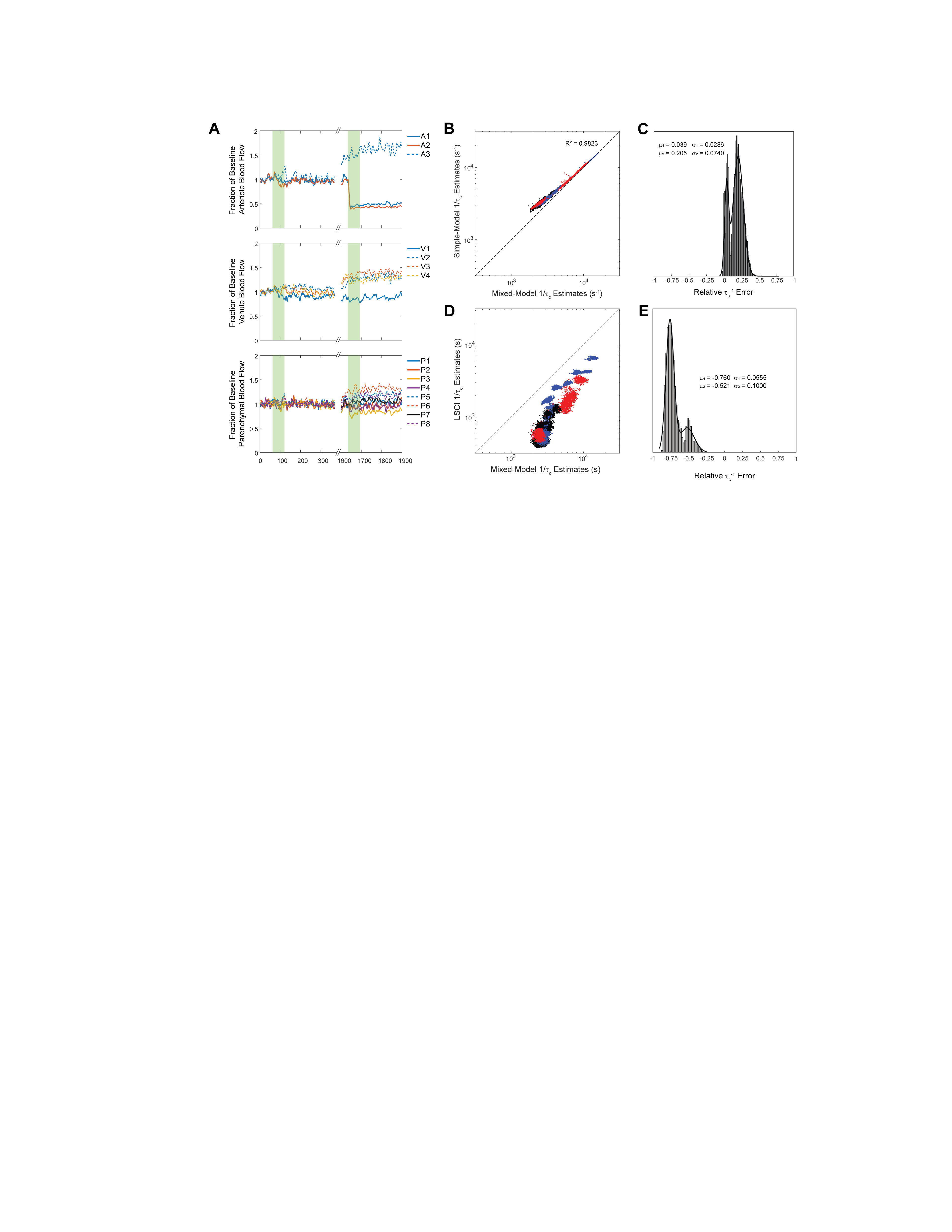

Figure S5. Correlation measures from simple-model fitting and LSCI deviate from mixed-model fitting during photothrombotic stroke. (A) Relative changes in the estimated inverse correlation times as estimated using the simple-model least-squares fitting approach for perfusion in arterioles (top), venules (middle), and parenchymal regions (bottom). (B) Scatter plot of inverse correlation time measurements estimated from ROIs in arterioles (red), venules (blue), and parenchymal regions (black) during photothrombotic stroke between simple-model fitting and mixed-model fitting as ground truth. While correlation is high, the points begin to deviate from the *y = x* dashed line at slower flows. (C) Distribution of relative errors of simple-model fitting to mixed-model least-squares fitting measurements with a fitted bimodal gaussian distribution. (D) Scatter plot of inverse correlation time measurements estimated for from ROIs in arterioles (red), venules (blue), and parenchymal regions (black) during photothrombotic stroke between single-exposure LSCI and mixed-model fitting as ground truth. Correlation between the methods is poor, with LSCI consistently underestimating flow. (E) Distribution of relative error between correlation times measured by LSCI and mixed-model fitting with a fitted bimodal gaussian distribution.

**Supplementary Movie 1.** sREMI output during mild photothrombotic stroke at 1-mW exposure. Green square indicates the illumination of the target arteriole by the green diode laser.

**Supplementary Movie 2.** sREMI output during moderate photothrombotic stroke at 2-mW exposure. Green square indicates the illumination of the target arteriole by the green diode laser.
